# cpiVAE: Robust and Interpretable Cross-Platform Proteomics Imputation

**DOI:** 10.1101/2025.11.20.689433

**Authors:** Yuxiang Li, ThuyVy Duong, Mary R. Rooney, Christie M. Ballantyne, Josef Coresh, Jennifer A. Brody, Nona Sotoodehnia, Dan E. Arking, Joel S. Bader

**Affiliations:** Department of Biomedical Engineering, Johns Hopkins University, Baltimore, Maryland, USA; Institute for Computational Medicine, Johns Hopkins University, Baltimore, Maryland, USA; Department of Genetic Medicine, Johns Hopkins University School of Medicine, Baltimore, Maryland, USA; Welch Center for Prevention, Epidemiology, and Clinical Research, Department of Epidemiology, Johns Hopkins Bloomberg School of Public Health, Baltimore, Maryland, USA; Section of Cardiovascular Research, The Texas Heart Institute at Baylor College of Medicine, Houston, Texas, USA; Optimal Aging Institute, NYU Grossman School of Medicine, New York, New York, USA; Cardiovascular Health Research Unit, Department of Medicine, University of Washington, Seattle, Washington, USA

## Abstract

Large-scale plasma proteomic studies often use different high-throughput affinity platforms, and measurements of the same protein across platforms are often discordant. Discordance hinders cross-study integration. Improving proteomics data integration would enable more powerful meta-analyses, improve statistical power for biomarker discovery, and provide a better understanding of proteome–phenotype relationships. Here we present a cross-platform proteomics imputation variational autoencoder (cpiVAE), a deep generative model for bidirectional imputation of protein abundances between two widely used platforms: Olink and SomaScan. Using a training cohort of paired measurements from the China Kadoorie Biobank (CKB), cpiVAE learns a joint latent representation that enables cross-platform imputation. The cpiVAE method improves benchmarks provided by established methods, k-nearest neighbors (KNN) Weighted Nearest Neighbors (WNN, from Seurat v4). The cpiVAE method achieves up to 30% higher correlation between imputed and true values than KNN and WNN. The cpiVAE method also generalizes well to an independent cohort from the Atherosclerosis Risk in Communities Study (ARIC). Without retraining, cpiVAE maintains high performance compared to benchmarks. Associations of imputed protein levels with clinical phenotypes closely mirror results using the actual measurements and increases power in a meta-analysis scenario. A post-hoc feature importance matrix enables interpretation of this AI model. Protein pair features extracted from cpiVAE have significant overlap with known associations in the Search Tool for the Retrieval of Interacting Genes (STRING) database. In summary, cpiVAE offers an accurate, generalizable, and interpretable solution for cross-platform proteomic imputation, enabling integrated analyses across studies with proteomics measured on different platforms. This user-friendly framework and pre-trained model weights are available under a BSD2 open source license at https://github.com/joelbaderlab/cpiVAE_v1.

## 1 Introduction

High-throughput affinity-based proteomics platforms have revolutionized large-scale biomarker discovery and genetic association studies [1, 2]. Two prominent commercially available technologies, Olink proximity extension assays (Olink) and SomaScan aptamer-based assays (SomaScan), each measures thousands of plasma proteins with high throughput [3–5]. These platforms differ in their biochemical detection methods, paired antibodies (Olink) versus single-stranded DNA aptamers (SomaScan) [6].

Each platform has advantages and disadvantages arising from the technical differences in detection methods due. Olink’s dual-antibody approach provides high specificity that in previous studies was thought to yield a greater number of protein-phenotype associations, whereas SomaScan’s single-stranded DNA aptamers offer broader proteome coverage and lower technical variability [7]. These differences mean that the same protein’s measured levels can vary substantially between platforms; furthermore, probes attributed to the same protein on different platforms may measure different isoforms or have additional contributions from closely related proteins. Thus, protein measurements from Olink and SomaScan are often weakly correlated. Multiple cross-platform studies of overlapping proteins have reported median Pearson correlations around 0.3, with substantial variability from protein to protein [7–11]. Thus, a simple approach of only aggregating signal for the same nominal protein across platforms has not worked well.

Additional technical factors underlie the limited concordance. High-abundance proteins tend to show better agreement between platforms [6, 9]. Low-abundance proteins or those with many values flagged as low quality often have poor cross-platform correlation. Platform-specific normalization and technical artifact can also contribute. For instance, SomaScan applies an adaptive normalization that, if not handled consistently, shifts its measurements distribution relative to Olink [12]. Additionally, each platform may target different isoforms or epitopes of the same protein. An aptamer might bind a different part or form of a protein than the antibodies. As a result, the same protein can have vastly different measurements.

These fundamental differences in measurements have hindered replication of associations and biomarkers across platforms [6, 7]. Prior analyses of matched Olink and SomaScan data have found distinct sets of significant protein–phenotype associations and protein quantitative trait loci (pQTL). In the CKB cohort, 2168 proteins were measured on both Olink and SomaScan, yet each platform uniquely identified certain signals [11]. Specifically, among 1694 proteins with one-to-one matching reagents, Olink detected cis-pQTLs for 765 proteins while SomaScan did so for 513, with only 30.2% reported as colocalizing. Similarly, one platform detected unique associations with clinical traits that the other platform did not [13]. This partial overlap suggests that combining data across platforms could provide a more complete picture of proteome–phenotype relationships. Indeed, an integrative analysis discovered additional health-related protein signals using both Olink and SomaScan, which would have been missed by either platform alone [14].

Variational autoencoders (VAEs) have been promising deep learning approaches for imputation [15]. VAEs learn a low-dimensional latent space of limited dimensionality using variational principles to enhance regularization and avoid overfitting. VAE architectures have performed well for integration of DNA and RNA genomics data, as seen in single-cell multi-modal studies [15]. Previous studies have also shown that VAE architecture can be problem-specific, and their advantage over simpler methods is not guaranteed in every setting, particularly if training data is limited. Recent benchmarks on external test sets have found that VAEs can under-perform simpler methods like k-nearest neighbors (KNN), or the related WNN implemented in Seurat V4 for genomic data [16–18]. Likewise, an importance-weighted autoencoder ensemble was applied to imputation between two metabolomics data platforms and was able to successfully recover the associations with clinical phenotypes [19].

Cross-platform imputation is related to the problem of imputing missing values for a single platform. Deep learning methods have been developed that improve on previous missing value imputation methods [20, 21]. VAEs and related architectures have been applied to denoise data and impute missing values in mass-spectrometry proteomics (distinct from the Olink and SomaScan platforms considered here). Autoencoder-based imputation of missing mass-spec measurements improved the detection of disease-associated proteins [22]. These advances suggest that, with enough training data and careful model design, deep generative models can learn the underlying relationship between multiple measurement techniques while mitigating platform-specific noise.

These previous studies motivate our exploration of VAE deep learning for cross-platform imputation between Olink and SomaScan. Our method, cpiVAE, performs cross-platform proteomic imputation using a joint variational autoencoder. We evaluate the accuracy of cpiVAE and its use in downstream applications including protein-phenotypic associations and protein quantitative trait loci (pQTL). Benchmarks are provided by KNN as a robust alternative and WNN as a state-of-the-art improvement over KNN architecture.

A crucial data resource is access to a cohort with the same samples measured on multiple platforms. We use data from the China Kadoorie Biobank (CKB), which generated proteomics data for thousands of individuals using by both Olink and SomaScan (Fig. 1) [11]. With CKB as training data, cpiVAE learns a joint low-dimensional representation of proteomic profiles that is shared between platforms. Because the encoding projects both platforms’ profile into a shared latent space, the method can decode from the latent space to predict either profile and impute cross-platform.

**Fig 1.**
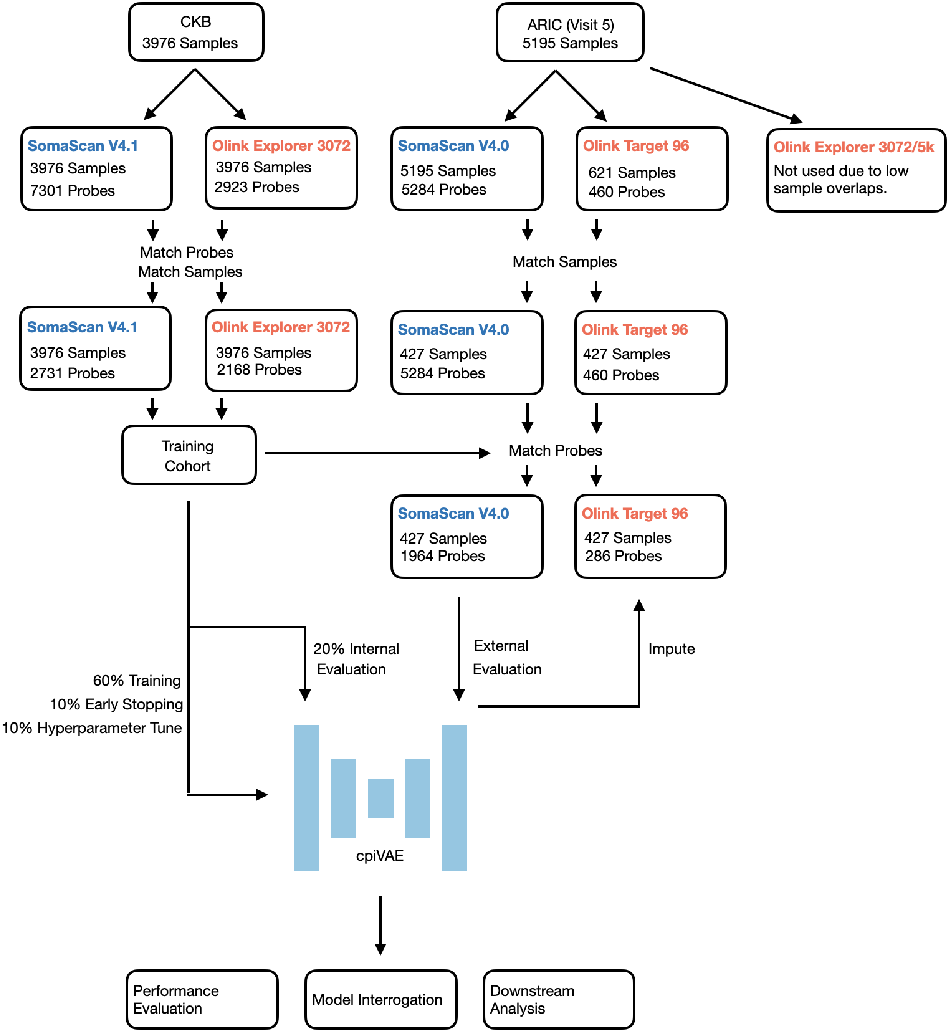
Overview of study design. The cpiVAE training and testing on CKB cohort and external evaluation on ARIC cohort. CKB provides paired Olink Explorer 3072 and SomaScan V4.1 data. ARIC contributes SomaScan V4.0 and Olink Explorer 96 data. Model training used 80% of CKB samples, testing using the remaining 20% of CKB, and external evaluation applied the CKB-trained model for SomaScan-to-Olink imputation for ARIC.

For technical evaluations, we assessed performance of cpiVAE verus KNN and WNN on held-out CKB data. Performance included imputation error and also correlation structure. For a practical assessment, we applied the cpiVAE learned from CKB, with no retraining or fine-tuning, to an independent cohort, ARIC [23]. The ARIC cohort is also demographically distinct from CKB, with very different genetic ancestry. We assessed the power to detect protein–phenotype associations using imputed versus measured data, and similarly compared cpiVAE with KNN and WNN. Finally, we explored model interpretability by interrogating the cpiVAE’s learned feature importance. Important features were extracted as protein–protein pairs, and these pairs were compared with known protein interactions and associations. We show that pairs that cpiVAE learns as important significantly overlap known protein–protein interaction (PPI) networks from the STRING database [24].

We provide cpiVAE as an open-source, user-friendly framework, enabling researchers to integrate and harmonize existing proteomic cohorts without requiring deep learning expertise. This approach facilitates the in silico expansion of datasets, a strategy analogous to how genotype imputation became instrumental in powering large-scale meta-analyses and accelerating discovery in genome-wide association studies (GWAS) [25]. Our framework thus overcomes a critical analytical barrier, empowering the proteomics and epidemiology communities to conduct more powerful, integrated studies across different experimental platforms.

## 2 Methods

### 2.1 Dataset and Study Design

We developed and trained cpiVAE using a multi-cohort design (Fig. 1). The term “probe” is often used rather than “protein” because in some cases, multiple assay probes may target different isoforms of the same protein, and conversely a single probe might recognize more than one protein. Our analysis treats each assay probe measurement as a distinct feature.

The primary training and internal validation data were from the publicly available China Kadoorie Biobank (CKB) proteomics sub-study. All participants were 30–79 years old at recruitment, drawn from 10 regions in China. In this cohort, 3976 adult participants had plasma samples profiled in parallel by both the Olink Explore platform and the SomaScan platform, yielding two complementary sets of 2731 and 2168 measurements for 2168 protein targets. These assays represent the overlapping set of proteins available on both platforms, measured under standardized protocols (Supp.

File 2, 3). The CKB dataset was randomly split into a 60% training set, a 10% early stopping set to prevent overfitting on model parameters, a 10% tuning set to prevent overfitting of model hyperparameters, and a 20% hold-out set for final internal performance evaluation. Imputation was bidirectional to explore the bilateral relationships between platforms.

The cpiVAE model trained on CKB was used without any further fine-tuning for assessment with an independent data set. This independent validation data came from visit 5 proteomics measurements from the Atherosclerosis Risk in Communities (ARIC) study, a longitudinal cohort of adults (aged 66-90 at the time of visit) from the United States. The ARIC proteomics dataset (Fig. 1) comprises 427 samples assayed on both platforms, along with corresponding clinical phenotypes and genotypes. Samples run on both platforms were typically analyzed using the Olink 96 panel, with only 286 probes. The Olink 3072 and Olink 5K platforms were not used due to few overlapping samples with the SomaScan platform. Therefore, for ARIC only imputation in the SomaScan to Olink direction was possible. Nevertheless, independent CKB and ARIC cohorts permitted evaluation of model generalizability under different ancestry compositions, environmental factors, and measurement sites.

Different versions of Olink and SomaScan datasets were used by CKB and ARIC, adding to the imputation challenge. Olink 3072 was used in CKB cohort, while Olink 96 data from ARIC cohort was used because of the large number of matched samples. Olink Explore 3072 is a modular library of eight 384-plex panels with Normalized Protein eXpression (NPX) output and low sample requirements. Olink Target 96 assays use 92 probes per panel and report NPX values derived from microfluidic qPCR Ct signals. Five panels were used in ARIC Olink 96: Olink CARDIOMETABOLIC, Olink CARDIOVASCULAR II, Olink CARDIOVASCULAR III, Olink INFLAMMATION, and Olink ORGAN DAMAGE. SomaScan V4.1 and V4.0 was used in CKB and ARIC cohort, respectively. All v4.x assays share the same core workflow and data standardization pipeline: hybridization control normalization, intra-plate median signal normalization, plate scaling, SOMAmer level calibration, and optional adaptive normalization (which was not used in this study). They differ in the number of probes, where v4.0 profiles 4,979 SOMAmer reagents while v4.1 expands the content to 7,596 total measurements.

### 2.2 Data Pre-Processing

Published post-QC CKB datasets were downloaded raw text tables from the FigShare data repository [26]. The original publication reported that all proteomic measurements had consistent quality control and normalization steps prior to public release. For Olink data, protein levels were reported as normalized using inter-plate controls and transformed using a predetermined correction factor. They were provided as NPX values on a log_2_ scale. The raw SomaScan data had been standardized using control samples and were reported as relative fluorescence units (RFU) for download. An additional adaptive normalization (ANML) step during the SomaScan v4.x standard protocol was omitted in both the CKB and ARIC datasets for better alignment of SomaScan and Olink data distributions. The ANML procedure, while useful in some contexts, can introduce shifts that exaggerate platform differences if not applied identically across datasets. After downloading, we applied log_2_ transformation to SomaScan before further analysis. Finally, we performed an additional feature-wise Z-score standardization. For each protein, values were scaled to mean 0 and standard deviation 1. This puts paired measurements on a comparable scale and helps the VAE training by roughly equalizing variance across inputs.

ARIC data was obtained from the ARIC Coordinating Center, and post-QC version datasets (denoted ‘cleaned’) were used for both Olink and SomaScan. For Olink, four internal controls were used. Sample plates with standard deviation above 0.2 NPX were excluded, and any sample that deviated 0.3 NPX from the median of the control was also excluded, resulting in 25 samples excluded in total. The ARIC Olink data included 4 samples with 124 missing values, which were dropped during evaluation. All other datasets have no missing values. We applied log_2_ transformation and feature-wise standardization to ARIC Olink data prior to further analysis.

For ARIC SomaScan, the median-normalized version datasets were used without ANML. Additionally, the manufacturer and ARIC-central QC provides optional quality flags for each aptamer that may indicate reliability issues such as non-specific binding or technical variability. We retained all probes regardless of their flag status to avoid inflating model performance or introducing bias by disproportionately removing low-abundance or highly variable proteins. Before probe matching, ARIC SomaScan v4.0 data included 879 probes with flag1, 329 probes with flag2, and 2620 probes with flag 3 (Supp. Table 3). Any values that fell below the assay’s limit of detection (LOD) were not removed from these datasets and remain as reported values. All features were mean standardized using parameters derived from ARIC’s own distribution to avoid any dependency on the CKB-derived scale.

### 2.3 Model Architecture

The cpiVAE model is a variational autoencoder designed for two data modalities (here, two proteomic platforms, although the framework supports any modalities and could be generalized to more than two platforms) (Fig. 2). During training, we explored a variety of more complex VAEs. Training takes multiple stages and because their accuracies were considerably lower in the initial stages, we focused on the simplest architecture. The cpiVAE model has two encoder networks that map each input profile into a shared latent space, and two decoder networks that map latent variables back to each platform’s data space.

**Fig 2.**
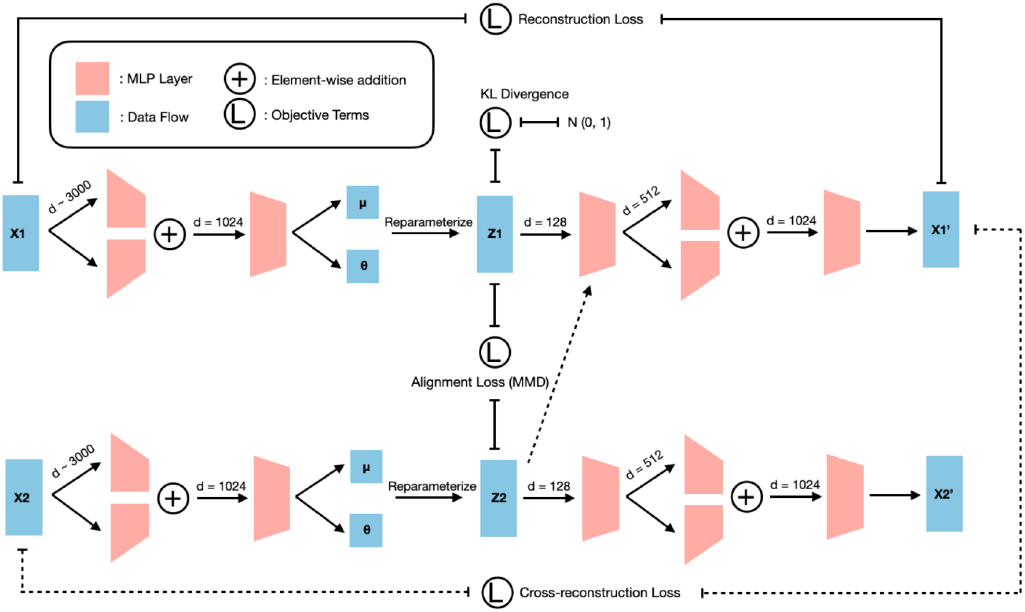
Schematic of the cpiVAE model architecture. The model comprises dual encoders (one for each platform) that map probe-level proteomics data through hidden layers to a shared latent space. The latent representations are regularized through Kullback–Leibler terms and aligned via Maximum Mean Discrepancy loss [27]. Dual decoders reconstruct both platforms from the latent space, enabling cross-reconstruction for imputation. Loss functions include reconstruction loss, Kullback–Leibler divergence, alignment loss, and cross-reconstruction loss. The architecture enables bidirectional imputation between platforms while learning biologically meaningful latent representations.

Given a dataset of paired observations 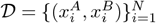 with 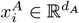 and 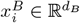, we use *•* ∈ *{A, B}* to denote the modality placeholder. We write 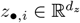 for the latent-space representation of sample *i*; *I* is the *d*_*z*_ *d*_*z*_ identity; and denotes elementwise (Hadamard) product.

Each encoder produces the parameters of a Gaussian posterior with diagonal covariance:

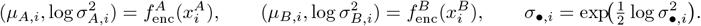

Latent samples are obtained with the reparameterization trick:

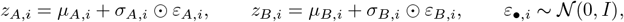

where the *ε*_*•*,*i*_ are independent across samples *i* and modalities *•*.

Self-reconstruction of each platform is enforced with an ℓ_2_ loss:

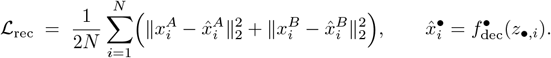

This corresponds to the negative log-likelihood under an isotropic Gaussian observation model with unit variance for each modality.

We then add an additional loss where each latent code is also decoded by the other platform’s decoder, to enforce transferable representations:

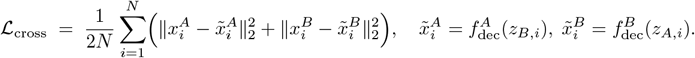

Each posterior is pushed towards the unit Gaussian prior with KL–divergence regularization:

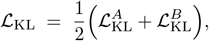

where

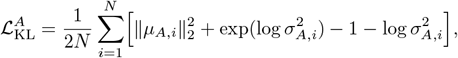

and analogously for 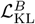. Here 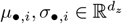; exp and log act element-wise, and the bracketed term is implicitly summed over latent dimensions.

To align the two posteriors we employ latent alignment as Maximum Mean Discrepancy (MMD) with an RBF kernel *k*_*γ*_ [27]. Specifically, we align the two *aggregated posteriors*

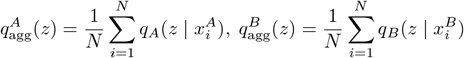

by computing 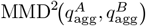. We estimate this on a minibatch using one Monte Carlo sample per item,

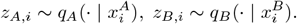

The resulting unbiased U-statistic estimator is

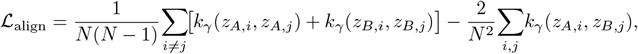

where

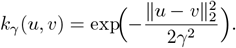

In practice, *γ* is chosen via the median heuristic on each minibatch.

The overall objective is composed of weighting the four contributions with non-negative coefficients *λ*_*{*rec,KL,align,cross*}*_ yields the total loss optimized during training:

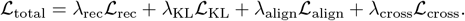

We empirically tuned the weights *λ* and other hyperparameters on a dedicated subset of the training data to balance the loss terms. In the final model, the trained values were *λ*rec = 0.90, *λ*_cross_ = 1.4, *λ*_KL_ = 1.4 *×*10^−4^, and *λ*_align_ = 1.9. These values reflect a balance between accurate reconstruction and cross-imputation and constraints that align the latent distributions aligned and provide just enough regularization to generalize. The KL term is scaled down to prevent over-regularization since each sample contributes 2168 reconstruction errors but on only 128 latent dimensions, effectively a 20*×* compression ratio. The relatively high alignment term enforces tight coupling of latent space representations. Each encoder/decoder network is a three layer resMLP. The middle layer is composed of a dual head residue block. ReLU activation functions, dropout layers, L2 regularization, and batch normalization are integrated into the network. All the network design and hyper-parameters are determined empirically on a split of tuning dataset using a Tree-structured Parzen Estimator (TPE). The cpiVAE framework generates an optimal configuration list upon automatic parameter searches. The full list of parameters and available tuning options are in Supplementary file (Supp.File 1). We initialized weights with Xavier/Glorot initialization and used the AdamW optimizer with learning rate 4 *×* 10^−4^ and mini-batches of 256 samples. Training proceeded for a maximum of 200 epochs, with early stopping applied if early stopping loss did not improve for 10 consecutive epochs. In practice, the model converged in under 100 epochs. Once the best hyperparameters were found, the model was retrained from scratch and evaluated on the 20% CKB internal test set and the ARIC external test set.

The cpiVAE code and trained model weights are available under an BSD 2 open-source license at our GitHub repository https://github.com/joelbaderlab/cpiVAE_v1. A release was also deposited on Zenodo 10.5281/zenodo.17245405. This repository also includes Python implementations of other VAE architectures, basic implementations of the KNN and WNN methods used as benchmark comparisons, preprocessing code, a Python-based pipeline to apply the trained model to new data, a BASH script to generate the results presented here, and additional documentation and examples. The model was implemented in PyTorch v2.7.1 with autograd and GPU acceleration. Model training and parameter search used a single NVIDIA A100 GPU. The model is lightweight with only 5 million parameters, compatible with consumer-grade GPUs or CPUs. We profiled resource usage against WNN and KNN methods on a consumer MacBook with M1 processing unit and 64Gb unified memory (see Results). Profiling was performed in part with Scalene [28]. Without computing feature importance, cpiVAE imputation inference on a 796 sample test dataset took 9.337 sec compute time, 62 MB disk space, 317 MB peak system memory, and 4 GB peak GPU memory (Supp. Table 1). Our WNN implementation with 5-fold validation took 39.703 sec compute time, 308 MB disk space, 1 GB peak memory and 2 GB peak GPU memory.

### 2.4 Methods for Benchmarking

We compared cpiVAE against two benchmarks, k-Nearest-Neighbor (KNN) and Weighted Nearest Neighbor (WNN). The KNN imputation imputed the target platform value as the weighted average of *k* measured values in the training data using Gaussian kernels. The value of *k* and kernel width were determined using a 5-fold cross-validation, with *k* = 15 for Olink → SomaScan imputation and *k* = 30 for SomaScan → Olink.

The WNN framework was introduced in Seurat V4 [16]. The WNN method learns sample-specific modality weights that quantify the relative information content of each data type for defining that particular sample’s state. Weights are learned by assessing the predictive power of each modality’s local neighborhood. In our application, WNN evaluates how well neighbors in the Olink space can predict a sample’s profile in the SomaScan space, and vice-versa. The resulting weights are used to construct a single WNN graph providing a unified view of sample-to-sample similarity. Imputation is enabled by supervised Principal Component Analysis (sPCA), which uses the WNN graph to find an optimal, low-dimensional projection of each modality that best captures the integrated cellular states. We adapted and extended the pyWNN codebase to implement an imputation pipeline following the sPCA-based reference mapping [16] and its corresponding Seurat v4 R implementation [29]. This multi-step process is as follows:

1. Perform a standard PCA on the scaled training data from both platforms.
2. Using these initial PCA embeddings, construct a WNN graph to learn the sample-specific modality weights and establish an integrated similarity metric between all samples in the training set.
3. Use the WNN graph as a smoothing operator on the original, high-dimensional training data. By performing PCA on this WNN-smoothed data, derive an sPCA transformation for each platform. This sPCA space is therefore optimized to emphasize features that are most consistent with the joint data structure.
4. Finally, train a Gaussian Kernel Regressor (KNN-based regressor with Gaussian kernel weight neighbors) to perform the final imputation. This regressor learns to predict the target platform’s values from the source platform’s representation in the learned sPCA space.

The WNN used 100 principal components for both the initial and supervised PCA transformations. The WNN graph construction began by identifying a union of 400 neighbors from each modality, which was then refined to a core graph of 40 neighbors per sample, using a small regularization term of 10^−5^ for numerical stability. Finally, the Gaussian Kernel Regressor was trained to perform imputation by weighting the 50 nearest neighbors in the learned sPCA space. Parameter values were selected using 5-fold cross-validation on a coarse grid (Supp. Table 2).

We constructed a negative control by permuting the sample-to-data mapping for modalities (equivalent to permuting each sample’s output vector), which maintains the covariance structure of each modality while making the modalities statistically independent. All methods were applied to the same training-test-evaluation splits in CKB for an unbiased comparison of performance metrics.

### 2.5 Performance Evaluation Metrics

Imputation accuracy was assessed per-protein and per-sample using the Pearson correlation coefficient *r*. In the CKB internal validation, for each protein with a nominal one-to-one correspondence of Olink and SomaScan probes, an *r* was calculated and summarized as a distribution of *r* values across protein features. Similarly, for each individual in the validation set, *r* values were calculated between imputed and actual imputed protein profiles to yield a distribution of sample-wise *r*.

External validation in ARIC consisted of two major analyses: phenotype association preservation and cis-pQTL recovery. For phenotype associations, we adjusted phenotypes for sex and age, performed association tests with the measured data, then selected the 3 binary and 3 continuous phenotypes with the largest number of significant association at 5% FDR. We then performed association tests for these phenotypes using imputed data and quantified the concordance of the effect estimates by the mean absolute errors of coefficients. In the summary plot, similar analysis was done on all 147 available phenotypes, of which 31 binary and 39 continuous phenotypes have at least one significant associations on truth dataset.

For cis-pQTL evaluation, we used the available genome-wide SNP data in ARIC to identify cis-pQTL for actual and imputed proteomic data. Associations were tested between each protein and all genetic variants within *±* 500 kb of its encoding gene, using linear regression adjusted for population structure. We applied a *MAF <* 0.05 and 5% FDR cutoff across all protein-SNP tests, which corresponded to approximately *p <* 10^−6^ in our analysis after multiple testing corrections. We further locus-pruned this set of 2353 hits down to 77 by selecting the most significant hits within a *±* 250 kb window. We assessed the overlap of cis-pQTLs hits identified by measured versus imputed datasets and the agreement between estimated allele effect sizes. The cis-pQTL analysis was conducted using TensorQTL [30].

### 2.6 Interpretability Analysis

The cpiVAE architecture can be interrogated to discover the relationships it has learned between proteins. A post-hoc interpretability analysis revealed which input feature proteins were most influential in imputing each target protein. This analysis used the Captum implementation of DeepLIFT to generate an importance matrix *I*_*q*→*p*_, with large value indicating that protein *q*’s measurement on one platform is heavily used by the model to predict protein *p* on the other platform [31]. Separate importance matrices were generated for Olink-to-SomaScan and SomaScan-to-Olink, restricting attention to proteins with a one-to-one correspondence between platforms.

The diagonal terms (self-importance) and off-diagonal terms (cross-importance) of the importance matrix were analyzed separately. The self-feature importance *I*_*p*→*p*_ was compared with per-protein measurement correlation *r*. For comparing the cross-importance and measured correlation between ligand-receptor pairs, we took the maximum value from both directions, and included all proteins belonging to the same family or subtypes as defined by the shared prefix of that pair.

For the off-diagonal terms, we hypothesized that proteins with a high importance relationship might be enriched for protein-protein associations and protein complex co-membership, with the abundance of an individual member of the complex serving as a good proxy for the abundance of the entire complex. Therefore, we compared the off-diagonal elements of the importance matrix with known human protein associations from the STRING protein-protein association database [24], restricting the STRING data to include only the proteins present in the importance matrix. We additionally compared to a smaller subset with physical interactions evidences. Self-interactions were explicitly removed for this comparison. Precision-recall curves were computed by systematically varying 16 logarithmically spaced importance threshold from stringent to permissive, calculating at each threshold the precision (fraction of predicted edges that are true reference edges) and recall (fraction of true reference edges that are recovered). The area under the precision-recall curve (AUPRC) was calculated using trapezoidal rule numerical integration. Statistical significance was determined by a permutation test with 1,000 iterations. In each permutation, STRING membership labels were permuted while keeping the importance values fixed. Each permuted dataset yielded a complete PR curve and AUPRC, building an empirical null distribution. The p-value was computed as (*k* + 1)*/*(*n* + 1), where *k* is the number of permutations yielding AUPRC values greater than or equal to the observed value and *n* is the total number of permutations.

## 3 Results

### 3.1 Proteomics Data

Two independent proteomics cohorts were used to develop and validate cpiVAE (Fig. 1). The dataset for training was the China Kadoorie Biobank (CKB), which provided paired Olink Explore 3072 and SomaScan V4.1 measurements for 2,168 proteins across 3976 individuals. We tested bidirectional imputation (between Olink and SomaScan) on a held-out portion of the CKB samples.

For external validation, we tested the CKB-trained model’s generalizability on an independent dataset from the Atherosclerosis Risk in Communities (ARIC) study, with paired SomaScan V4.0 and Olink Target 96 data reported for 427 individuals. Due to the smaller Olink panel used in ARIC, evaluation was limited to unidirectional SomaScan-to-Olink imputation for the 286 proteins available on that panel (Table 1). This design allowed us to assess imputation performance on a cohort with different ancestry and assayed with different platform versions, providing a robust test of its real-world utility.

**Table 1.** Summary of proteomics cohorts used in the study.

| Cohort | Shared Samples | SomaScan Probes | Olink Probes | Shared Proteins |
| --- | --- | --- | --- | --- |
| CKB | 3,976 | 2,731 (v4.1) | 2,168 (Explore 3072) | 2,168 |
| ARIC | 427 | 1,964 (v4.0)* | 460 (Target 96) | 286† |

\*Number of probes used as input for evaluation.
†Number of probes used as ground truth for evaluation.

### 3.2 The cpiVAE method outperforms existing methods in cross-platform proteomics imputation

We developed cpiVAE, a joint variational autoencoder neural network architecture with dual encoder and decoder heads (Fig. 2). We used cpiVAE to impute between two high-throughput proteomic platforms, Olink (Thermo Fisher Scientific) and SomaScan (SomaLogic). We benchmarked cpiVAE against 2 baseline methods, KNN and WNN.

We trained cpiVAE, as well as KNN and WNN, using 80% of the CKB data, holding out 20% for testing (see Methods). We first quantified how well each method reconstructed held-out measurements in the CKB dataset. We calculated Pearson’s *r* between imputed and observed values for each of the 2168 shared probes for both imputation directions. While all are bi-modal, cpiVAE’s distribution is more sharply centered at a median of 0.609 for S → O and 0.488 for O → S, a noticeable 28.5% and 25.7% improvement compared to the next best method, WNN (medians of 0.474 and 0.381) and above the KNN (Fig. 3A, left). The negative control using permuted-VAE is centered near 0, confirming that any real signal yields substantially higher correlations. Aggregating across features within each individual also shows higher performance of cpiVAE, which attains median correlations of 0.520 and 0.477, outperforming KNN (0.411, 0.371) and WNN (0.358, 0.373, Fig. 3A, right).

**Fig 3.**
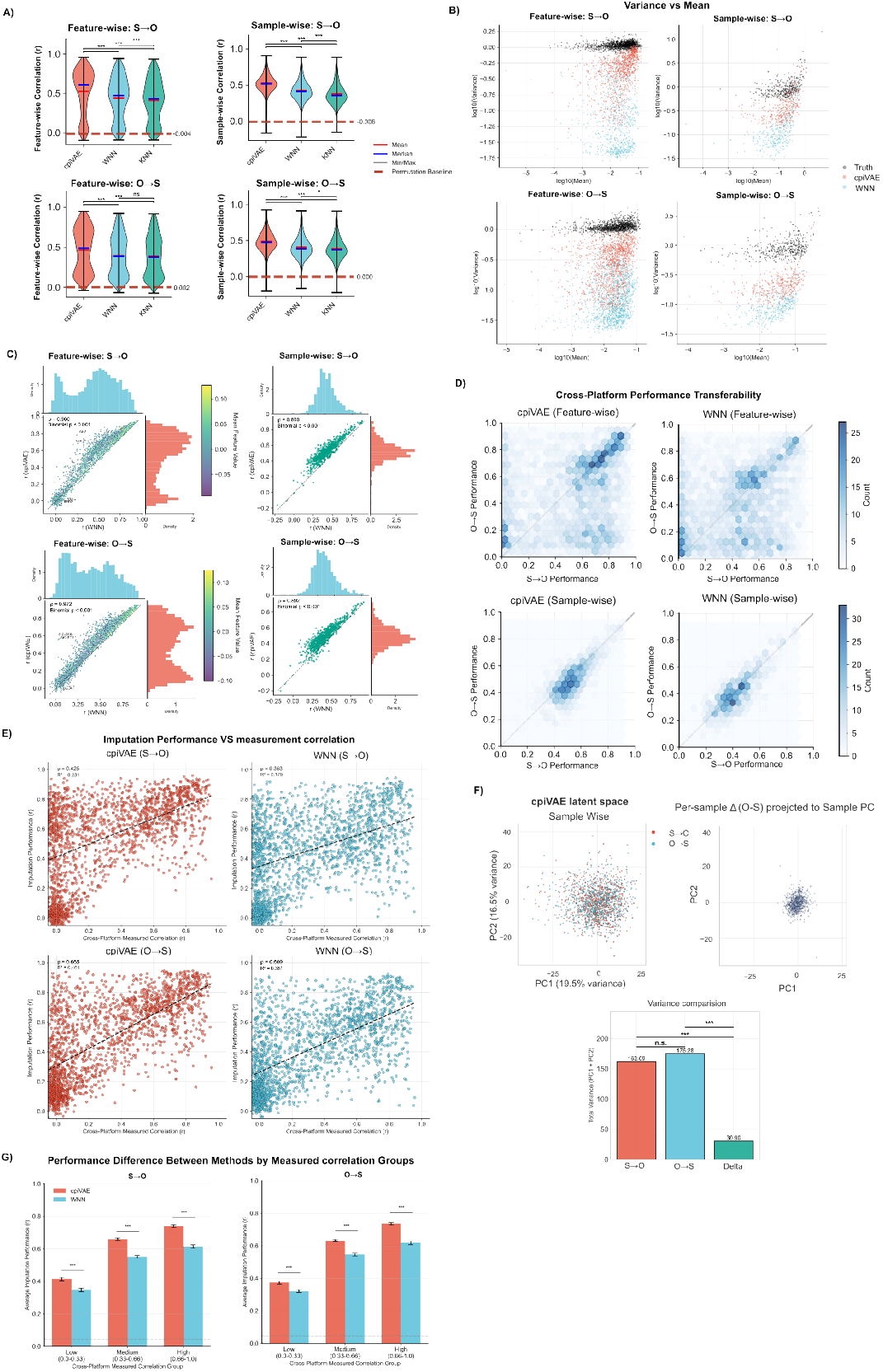
Cross-platform proteomics imputation performance comparison. (A) Violin plots of Pearson correlation *r* between imputed and measured values. Left: prob-wise distribution; Right: sample-wise distribution; Top: S → O; Bottom: SomaScan dataset. All the following plots follow the same arrangement. Horizontal bars indicate medians and means; whiskers show full range; red dotted lines indict mean correlation of permuted negative control; Mann-Whitney U test is used for statistical significance. (B) Mean versus variance plots for measured and imputed datasets. Both axes are presented in log scale. Color indicate methods. (C) Scatter plots with marginal histograms shows the cpiVAE accuracy compared to WNN. Dashed line indicate identity. Data point color reflects mean protein abundance. (D) Cross-platform consistency density plot: protein accuracy in the Olink to SomaScan direction is plotted against accuracy in the reverse direction. Dashed line shows identity. (E) Relationship between measured cross-platform correlation and cpiVAE imputation accuracy. Red dashed line indicate linear fit. (F) Top right: principal-component analysis of the cpiVAE latent space for samples from both datasets. Color indicate origin. Top left: the differences between two measurements of the same sample, projected to the principle components space of original space. Bottom: variance comparison between three data distributions. Levene test for significance. (G) Imputation performance comparison between methods grouped by measured cross-platform correlation. Mann-Whitney U test for signficance.

In addition to providing a better correlation, cpiVAE also generates distributions with variances that more closely match the true variance (Fig. 3B). These results suggest that KNN and WNN may over-regularize, shrinking estimates towards a mean. While cpiVAE also regularizes, the effect is not as large.

As expected, WNN consistently outperformed KNN in performance. Some comparisons below omit KNN to simplify the presentation. Overall, cpiVAE improves on WNN (and KNN) for the vast majority of proteins and individuals (Fig. 3C).

Correlations between imputed and measured values are better for cpiVAE than WNN for 84% of samples and 92% of probes, a substantial improvement (*p <* 0.001, two-tailed binomial test). The cpiVAE method performs better both for low-abundance proteins like ABO and high-abundance proteins like SIGLEC.

To assess whether imputation performance depends more on the protein itself than on the platform, we compared results across datasets in both O→ S and S → O directions for the 2168 proteins shared between platforms (Fig. 3D). The weak but significant correlation of cpiVAE *r* = 0.283 and WNN *r* = 0.299 indicates that many probes exhibit different performance depending on the direction of imputation. We observed three main clusters in the protein performance: (1) proteins that are imputed well in both direction; (2) proteins imputed well S → O but poorly from O →S; and (3) proteins imputed poorly S →O and slightly better imputation O →S. These differences may be due to differences in dynamic range or noise between platforms for these proteins. Our model does not eliminate those differences, which may not be possible to correct with any imputation method.

We also assessed whether performance of cpiVAE and WNN were primarily driven by proteins shared between Olink and SomaScan. For each protein present on both platforms, the correlation between Olink and SomaScan measured values itself correlates with cpiVAE imputation accuracy (Fig. 3E, left panels). Nevertheless, we also observed many proteins with near-zero measured correlation between platforms yet with good cpiVAE imputation performance (upper-left quadrant). The results from WNN are similar, except that there are a few more highly correlated proteins that are low performing (Fig. 3E right panel). Grouped by cross-platform measured correlation, cpiVAE imputation improved on WNN (Fig. 3G).

We finally investigated the 128-dimensional latent space of cpiVAE by projecting the latent space onto its first two principal components (this type of analysis is not possible for WNN and KNN, which lack a latent space). Projecting the shared latent space with PCA demonstrates that encodings of Olink and SomaScan measurements form a single cloud rather than two modality-specific clusters (Fig. 3F, top left). This overlap confirms that the alignment loss successfully joins both platforms onto a common manifold, a prerequisite for unbiased bidirectional decoding. We then projected the differences between matched samples on both platforms onto the same principle components. We observed a tighter distribution, showing that the between-platform variance is significantly smaller than between-sample variance (Fig. 3F, top right and bottom).

### 3.3 cpiVAE imputation generalizes to independent cohorts of different ancestry

The cpiVAE model trained using the CKB cohort (East Asian ancestry) was then applied directly to the ARIC cohort (self-identified European and African ancestry) for imputation from SomaScan to Olink to determine whether cpiVAE could generalize to an independent and demographically distinct cohort measured at a different facility. Furthermore, ARIC used a different SomaScan version than CKB and a different Olink platform of 96-plex panels, which profiled fewer proteins than the CKB Olink platform. These differences present a challenging scenario for generalization.

Across the 286 proteins that overlap the Olink 96 panels used in ARIC, cpiVAE produced the tightest distribution of feature-wise Pearson correlations with the measured values, achieving median *r* = 0.557 (a 42.10% increase from WNN, Fig. 4A). WNN and KNN had slightly lower medians and broader tails, and the permuted control centered near zero as expected. In per-sample assessments, cpiVAE again had the best performance with median *r* = 0.524; WNN and KNN had median *r* = 0.366 and 0.329, respectively, significantly worse than cpiVAE (*p <* 0.001, Mann-Whitney U test). The cpiVAE method had only minimal feature-wise and sample-wise generalization error assessed by performance drop-off between internal testing and external validation (Fig. 4B). In contrast, WNN had worse generalization error in sample-level accuracy (Δ*r* = 0.053).

**Fig 4.**
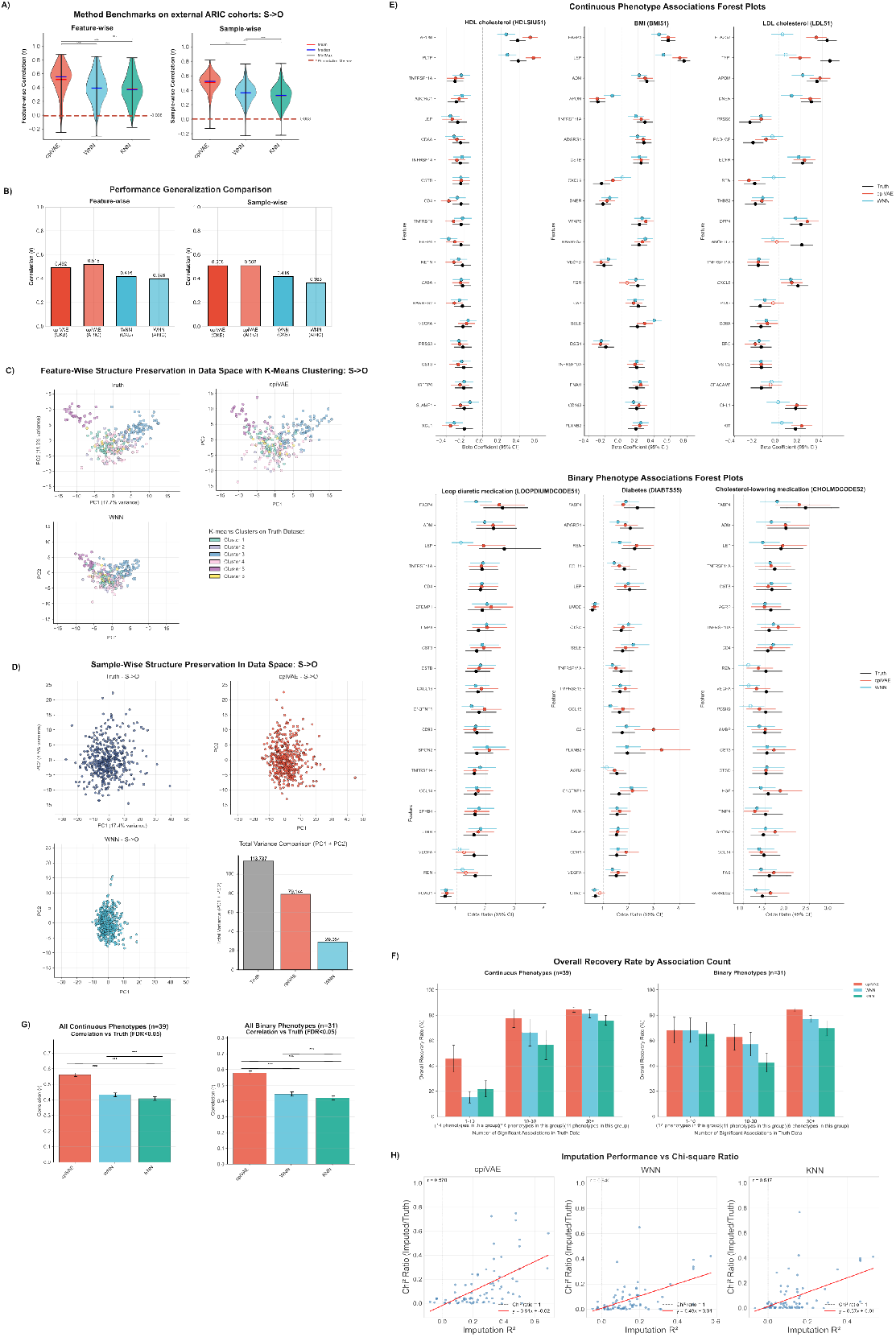
cpiVAE imputation generalizes to independent cohorts of different ancestry. (A) Violin plots of Pearson correlations between imputed and measured Olink values in ARIC, computed per feature (left) and per sample (right). Colors denote methods; horizontal bars indicate medians and means; whiskers show full range; red dotted lines indict mean correlation of permuted negative control; statistical significance from Mann-Whitney U. (B) Performance degradation from internal to external cohort testing. (C) Feature-wise PCA after coloring by k-means clusters (*k* = 6, clustered on measured dataset). The cpiVAE and WNN imputed datapoints are projected onto measured truth PCs. (D) PCA plot of measured and imputed sample-wise Olink profiles. Bottom right: sample-wise total variance comparison for truth and imputation dataset. (E) Forest plots for the quantitative and binary phenotypes with the highest numbers of significant hits, showing top 20 significantly associated proteins for each phenotype. Points represent size estimates, horizontal bars the 95% confidence intervals. Colors represent methods. Filled versus open circles indicate significance at 5% FDR. (F) Summary mean significant hit recovery rate between methods using all significant hits in all continuous (left) and binary (right) phenotypes at 5% FDR. Data is grouped by the number of significant associations in each phenotype. (G) Summary mean correlation to truth dataset comparison between methods using the same all FDR significant hits in all continuous (left) and binary (right) phenotypes. (H) Recovery of genetic association strength scales with imputation accuracy. For the n=2353 protein–variant pairs with significant cis-pQTL signal in the measured data (truth *p <* 10^−6^; MAF *>* 0.05), the ratio of the *χ*^2^ statistic from imputed data to that from measured data is plotted against per-protein imputation performance as *R*^2^. Dashed line marks perfect recovery; red lines marks least-squares fits.

The cpiVAE method also retained the correlation structure of the data better than WNN, indicated by PCA projections of features (Fig. 4C) and samples (Fig. 4D). Olink probes were clustered into 6 groups using *k*-means clustering of the measured value. PCA were then performed on measured truth dataset. Values imputed by cpiVAE and WNN were projected onto the truth PC space. With cpiVAE, imputed and measured values for each cluster generally co-localized in the PCA projection (Fig. 4C, top right). With WNN, the imputed values showed less variation (Fig. 4C, bottom left). This over-regularization by WNN was similarly noted for CKB data. A PCA projection of samples also shows that values imputed by cpiVAE have a distribution similar to the measured values, whereas values imputed by WNN have greater shrinkage (Fig. 4D).

We next assessed the use of imputed values for phenotypic association, with results presented for continuous and binary phenotypes with large numbers of significant associations in the actual measured truth dataset (Fig. 4E). Associations performed using values imputed by cpiVAE had effect sizes and confidence intervals close to the values obtained using the true data. Associations from data imputed using cpiVAE generally retained significance more often than WNN.

The recovery of significant associations was assessed systematically across all phenotypes, grouped by the number of significant associations using the actual data (Fig. 4F). The cpiVAE method gave the highest recovery rate, 46 out of 69 associations (66.9%). In comparison, data imputed by WNN only recovered 39 associations (57.3%), and KNN only recovered 36 associations (51.8%). As expected, permuted data yielded no significant associations.

The effect sizes, defined as regression coefficients for continuous phenotypes and log-odds ratios for binary traits, were compared for imputed and measured data. The cpiVAE method yielded the highest correlation for both log-odds ratios (*r* = 0.578) for binary traits and for beta coefficients (*r* = 0.560) for quantitative traits (Fig. 4G). Values for WNN were worse, *r* = 0.445 and *r* = 0.433, and KNN were worse still, *r* = 0.419 and *r* = 0.408. The cpiVAE correlations were significantly better for continuous and binary phenotypes (*p* = 8 × 10^−39^ and *p* = 1.4 × 10^−30^, Wilcoxon Signed-Rank Test).

Recovery of genetic associations was also assessed across imputation methods. Using measured data, 77 significant cis-pQTL protein–variant pairs were identified (MAF ≥ 0.05, FDR ≤ 0.05, locus-pruned to be most significant within ± 250 kb window). Because power is limited by the small cohort size, we used the *χ*^2^ test statistic for cis-pQTL to assess performance. The *χ*^2^ for imputed versus measured data increased monotonically with per-protein imputation *R*^2^ (Fig. 4H). This relationship was strongest for cpiVAE (Pearson *r* = 0.570), with a regression slope nearly twice that of KNN and substantially greater than WNN, indicating that improvements in imputation accuracy yield proportionally larger gains in recovered genetic signal. Although ratios remain below 1, as expected from attenuation due to measurement noise, cpiVAE consistently retained a larger fraction of the true test statistic than the benchmarks. The permuted control showed no relationship between imputation performance and genetic signals recovery (*r* = − 0.036). Thus the improved technical accuracy of cpiVAE should also improve the power to detect genetic associations with imputed data.

### 3.4 cpiVAE increases the power of meta-analysis

In ‘classic’ meta-analysis, a protein feature present on both platforms can be analyzed separately on each platform, and then *z*-scores or other test statistics may be combined. Imputation enables meta-analysis for a harmonized set of protein features, with different workflows possible, ‘imputation-separate’ and ‘imputation-combined’. The imputation-separate strategy treats the imputed dataset as a stand-alone dataset, similar to the classic approach: association metrics are computed separately for the native and imputed cohorts and then and combined. In contrast, the imputation-combined strategy merges the imputed samples onto one harmonized dataset, which is then analyzed as a single larger cohort.

To compare with ground truth and with the classic approach, our assessment focused on the 255 ARIC protein features subset that are present on both the Olink and SomaScan platforms. We randomly assigned samples to two sets to simulate a scenario in which cohorts are run on different platforms, repeating the splitting 20 times (Fig. 5A). We then performed imputation from SomaScan to Olink and performed meta-analysis using imputation-separate and imputation-combined workflows. A set of true associations was defined by analyzing the actual Olink data available for all the samples.

**Fig 5.**
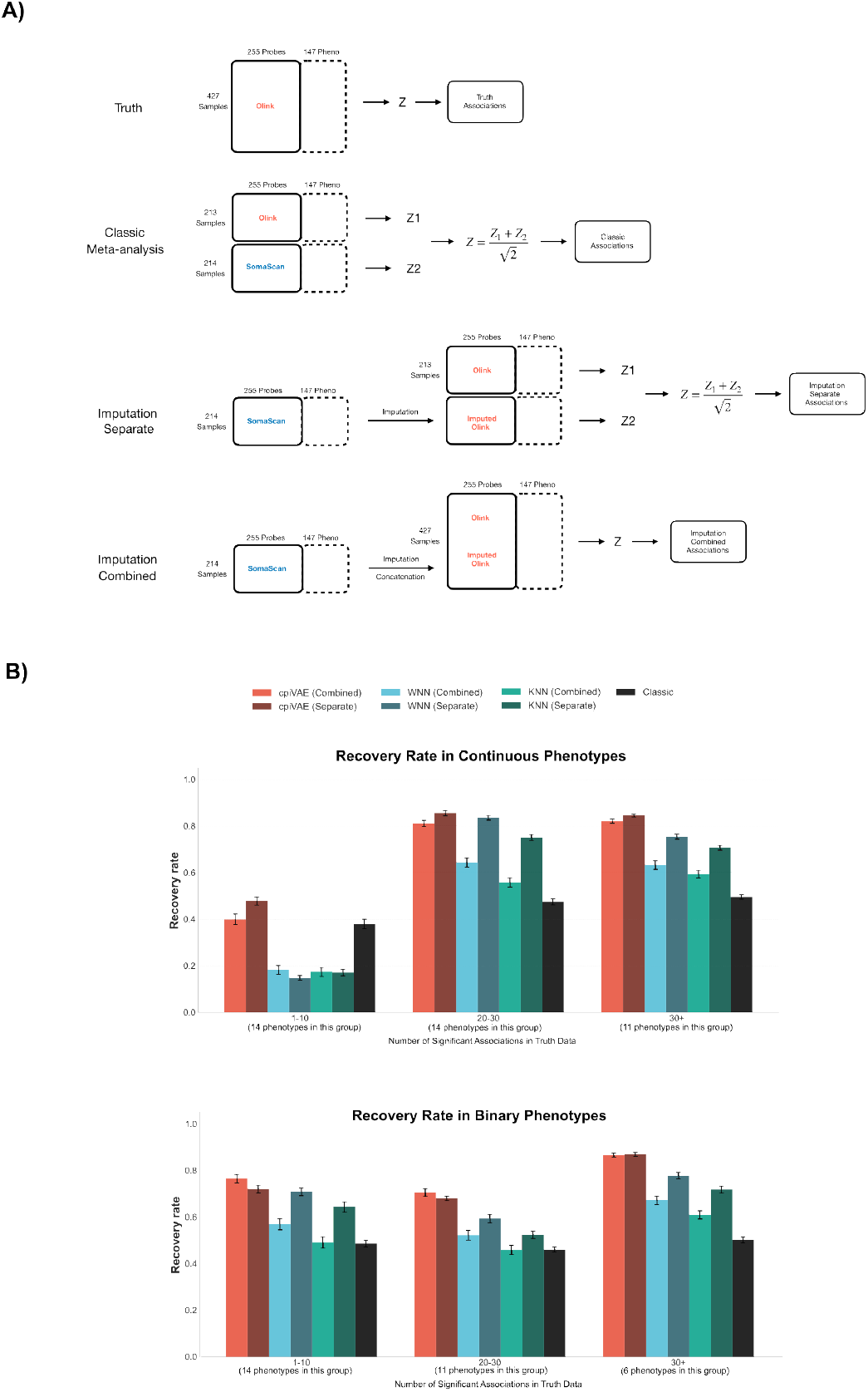
Imputation improves cross-platform meta-analysis and association discovery. (A) Schematic of 4 different meta-analysis procedures. (B) The performance of conventional meta-analysis, imputation-separate strategy, and imputation-combined strategy evaluated by the mean recovery rate of significant associations for all available phenotypes, stratified by the number of true associations for each phenotype. Error bars indicate standard deviations from 20 random assignment of samples into two groups.

Our results demonstrated considerable advantage when using any imputation methods compared to classic meta-analysis (Fig. 5B). The performance improvement increased for phenotypes grouped by number of protein associations. Of the different imputation methods, cpiVAE generally performed best, followed by WNN and then KNN. Imputation-separate (calculating *z*-scores separately for native and imputed data) generally performed better than imputation-combined (calculating *z*-scores from a single harmonized data set), and both were far superior to classic meta-analysis. Poor performance of classic meta-analysis may arise from protein features that nominally measure the same protein but actually report on different species.

Considering assessments of technical accuracy, phenotypic associations, pQTL identification, and meta-analysis, cpiVAE generalizes to external cohorts better than state-of-the-art methods. These results suggest that cpiVAE could be a standard approach for harmonized analysis of cohorts measured using Olink and SomaScan platforms.

### 3.5 cpiVAE model interpretation aligns with known protein associations and physical interactions

While model performance is an important criterion for imputation, an additional goal is model interpretation: can the model be investigated to understand what leads to superior performance, and do these factors provide additional insight? Importance scores, which define the contribution of an input feature to an output prediction, can be calculated for cpiVAE (see Methods). Importance scores provide insight because both the input features and the output predictions are proteins, and the scores can be compared with known protein associations and physical interactions. Protein-protein importance scores from cpiVAE were compared with known protein-protein associations and physical interactions.

The feature importance matrix was calculated using held-out CKB set (see Methods). Visualization of a randomly selected subset of *I*_*q*→*p*_ matrix for both directions indicates strong self-importance for many proteins as a bright series along the main diagonal (Fig. 6A). Self-importance is strongly associated with consistency of same-protein measurements across platforms, with significant correlation in both the Olink to Somascan (*r* = 0.70, *p* = 2.9 *×*10^−307^) and the Somascan to Olink (*r* = 0.73, *p* = 9.7 *×* 10^−302^) direction (Fig. 6B).

**Fig 6.**
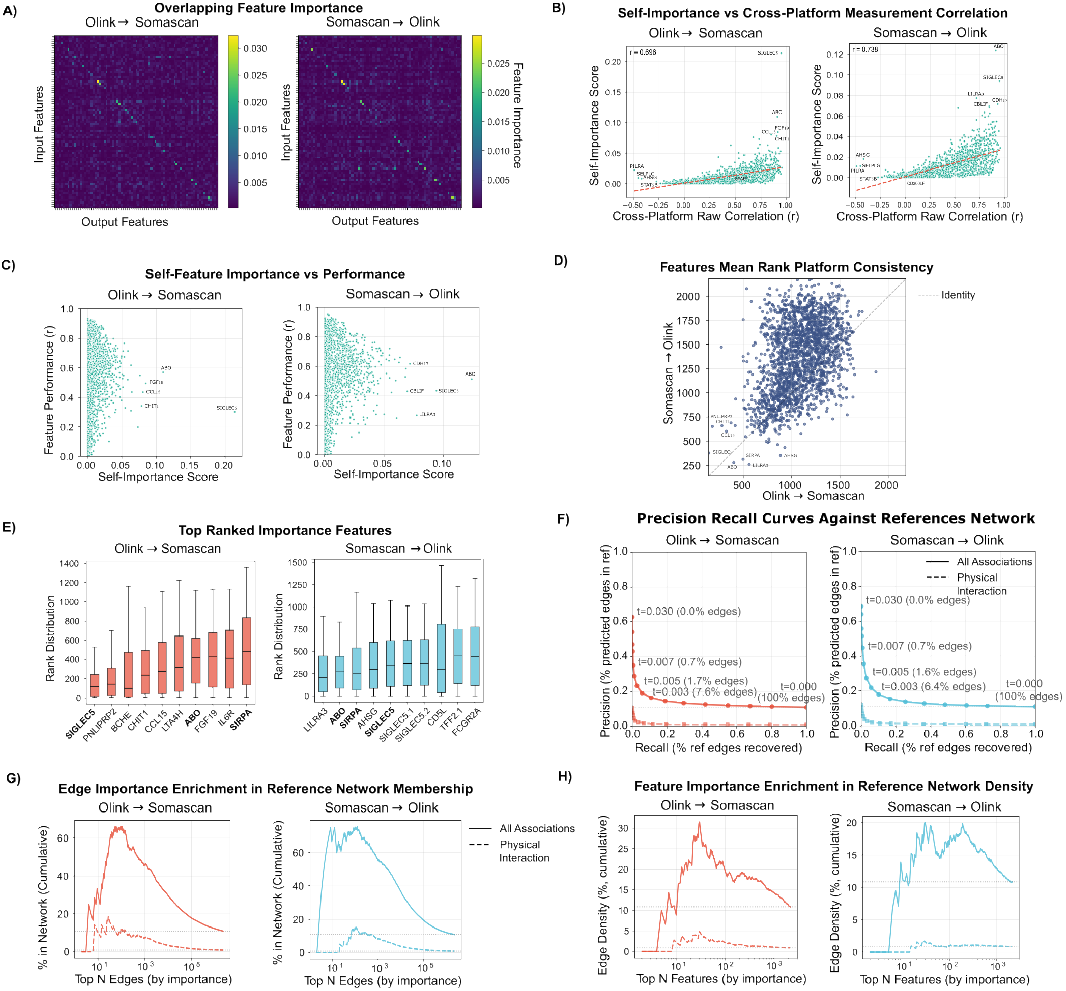
(A) Heat maps of the DeepLIFT importance matrix for Olink → SomaScan (left) and SomaScan → Olink (right). All the following subplots follow the same arrangement. Random subset of 100 features is shown. Color scale denotes importance score. (B) Relationship between a protein’s self-importance score and its cross-platform imputation accuracy *r*. Dashed lines shows linear fits, with *r* values reported in the panels. Selected outliers are annotated. (C) Relationship between a protein’s measured platform correlation and self importance score. Selected outliers are annotated. (D) Feature importance rank-cross platform consistency, representaed as the mean rank of each input feature among all output features. Dotted line denotes identity importance. (E) The 10 most important features used by the model, ordered by their mean ranks among all predicted features. (F) Precision–recall curves for recovery of STRING protein associations (solid line) and a physical interactions (dashed line) by cpiVAE protein-protein importance, with points labeled by importance thresholds. (G) Enrichment plot between the importance of edges and cumulative percentage of edges captured by the STRING associations (solid line) and physical interactions (dotted line). (H) Enrichment plot between the average importance of network feature nodes (proteins) and the cumulative edge density of the nodes.

High self-importance is not required for high imputation accuracy, however, as shown in a comparison of the self-importance score with the correlation *r* between actual and imputed measurements (Fig. 6C). The correlation is positive but weak (Olink to SomaScan, *r* = 0.31; SomaScan to Olink, *r* = 0.22), with many proteins imputed well despite lack of concordance between platforms. High self-importance is not sufficient for good imputation, however. Highlighted outliers with high self-importance but below-average imputation include SIGLEC5 and ABO (Fig. 6E).

The cpiVAE method can perform well even when direct measurements are not concordant because it can draw information from other proteins across the panel, corresponding to off-diagonal elements of the importance matrix (seen as off-diagonal bright elements in Fig. 6A). Cross-importance tends to involve the same proteins in the two imputation directions. For example, the cross-importance of protein pairs in the Olink-Somascan and Somascan-Olink direction is moderately correlated (Fig. 6D, *r* = 0.401, *p* = 8.2 *×* 10^−83^). And among ten proteins with the greatest overall importance, three are shared between networks (Fig. 6E, *p* = 4.5 *×* 10^−74^, Fisher exact test).

Many protein pairs with strong cross-importance correspond to known biology. For example, strong cross-importance is observed for TNFR (tumor necrosis factor receptor) with its ligand TNF (max importance = 0.028, top 0.01%); for IL-1R (interleukin-1 receptor) and its ligand cytokine IL-1 (max importance = 0.029, top 0.01%); and for LEPR and its ligand LEP (leptin) (importance = 0.018, top 0.03%). Cross-importance can be high even when direct correlations are low. The proteins EGF (Epidermal Growth Factor) and EGFR have high cross-importance (score = 0.015, top 0.05%) despite almost no correlation in the raw data (maximum absolute *r* = 0.014, top 69%).

To test the hypothesis that importance corresponds to biology systematically, protein pairs ranked by cross-importance were compared known protein-protein functional associations and protein-protein physical interactions obtained from the STRING database [24]. Precision-recall curves indicate strong enrichment of associations and interactions among protein pairs with high cross-importance (Fig. 6F, *p* = 0.001, see Methods). Protein pairs top-ranked by importance were compared with known associations or interactions (Fig. 6G). Enrichment was observed for the top 100–10000 pairs, corresponding to the top 0.004-0.4% of all edges. In addition to identifying important protein pairs, important individual proteins were identified by their total ranked importance in the imputation matrix. Known associations and interactions were enriched between proteins with high individual importance (Fig. 6H).

These results indicate that the cpiVAE model has learned real relationships between proteins, including both functional associations and physical interactions. Such relationships were not explicitly given to the model, yet emerged because exploiting them improved cross-platform prediction.

## 4 Discussion

The cpiVAE method uses a deep learning framework to harmonize and integrate data from different proteomics platforms, solving a practical problem for population proteomics. Imputation has been crucial for genetics by enabling meta-analysis across multiple smaller cohorts to have increased power. In proteomics, cross-platform imputation has been more challenging because different platforms are used, measuring different subsets of proteins, and even for shared protein features not necessarily measuring the same species. Additional complications are multiple versions of different panels within individual platforms. It is common in practice to pool shared proteomic results across studies. However, given the modest cross-platform correlations (median *r* ~ 0.3) observed between Olink and SomaScan, such naive integration can dilute true signals and introduce inconsistencies. By learning a shared representation of Olink and SomaScan proteomic platforms, cpiVAE can accurately impute one platform using data from the other. The cpiVAE method should enable an increase in power for past and future proteomics studies. Our implementation and trained weights are available under a permissive open source license.

Our results demonstrate that cpiVAE outperforms other imputation methods, generalizes effectively to independent cohorts, and learns a model structure that aligns with real biology. A primary advantage of cpiVAE is its ability to exploit complex multivariate structures in the data. Traditional methods like nearest-neighbor imputation or linear regression use limited local data similarity. As shown, this independence to the basis vectors of observed individuals is limited when cross-platform relationships are not straightforward. The cpiVAE, in contrast, can capture nonlinear and higher-order correlations among proteins, effectively learning how groups of proteins covary across individuals. This proved crucial for imputing proteins with weak direct correlations between platforms: the model “borrows strength” from biologically related proteins to infer their levels. As a result, cpiVAE provided substantial improved powers for those hard-to-predict proteins, which are often of great interest.

Moreover, generalizability has often been a struggle in deep learning approaches in biology due to limited number of samples. This is evidenced in the case of deep learning of single cell multi-modality integration, where mixed results were reported in external cohort benchmarking. In our testing, cpiVAE shows almost no performance degradation between a training cohort and a completely independent application cohort, with proteomics of entirely different populations and measurements conducted at separate facilities.

Additionally, cpiVAE is superior to state-of-the-art methods in preserving the true phenotypic associations signals when applied to new ARIC datasets. This suggests that the model learned the robust underlying biology rather than overfitting to cohort-specific noise. This likely reflects our stringent training pipeline, incorporation of regularization and the inherent noise robustness of the VAE approach.

Interestingly, in preliminary experiments we tried several more complex VAE variants and modern generative network components including vector-quantized VAE (VQ-VAE), variational mixture of posteriors (VampPrior VAE), importance-weighted autoencoders (IWAE), U-Net style architectures of various sizes, as well as attention mechanisms and even a hybrid VAE-GAN approach. However, the simpler cpiVAE architecture outperformed all these more complicated architectures (numerical results not reported because alternatives could not be completely trained). Any training data corpus has a tradeoff between complexity and performance. Because proteomics imputation has lower intrinsic complexity compared to image or language data, as well as a limited sample size, overly complex models were prone to overfitting or did not yield additional benefit. The chosen architecture provided a balance of expressiveness and regularization for our data.

The cpiVAE model is interpretable, with protein-protein importance scores that overlap with known protein-protein associations and physical interactions, including ligand-receptor interactions. Importance edges are significantly enriched for protein associations and physical interactions. By highlighting potential interactions or co-regulations not obvious from standard analyses, cpiVAE may be able to generate novel hypothesis. For example, many high-weight edges involved receptor–ligand pairs. In plasma, these often included soluble receptors. Our finding may reflect that ligands can circulate partly in receptor–ligand complexes. Olink and SomaScan may therefore differentially capture free versus complexed species, and the model’s cross-weights may reflect borrowing information from an interaction partner. Second, edges occurred within protein families, where closely related proteins share domains. While these may indicate co-regulation, they could also arise from partial cross-reactivity or non-specific binding of the probes. Accordingly, edges in the learned network are best interpreted as informative dependencies for prediction, with orthogonal tests helping to distinguish biological coupling from probe cross-reactivity.

From a practical perspective, cpiVAE can enable new analyses that were previously infeasible, such as cases where proteomic data from different studies or subcohorts cannot be directly combined due to platform differences. Cross-platform imputation can also give access to the union of proteins measured by either platform, rather than the intersection. Our goal in providing open source and weights is to advance these types of studies for the community.

The cpiVAE method has limitations. Where possible, measuring both platforms or a gold-standard like mass spectrometry remains ideal for cross-validation. It requires a reasonably large paired dataset for training. Aggregate multiple smaller paired studies may permit training of a more robust model. Additionally, while cpiVAE preserves many known signals, differences between platforms could be lost due to regularization. While our interpretability analysis was insightful, the feature importance scores are approximate and can be influenced by co-linearity. Alternative approaches including integrated gradients and gradient SHAP (both incorporated in our implementation) yielded qualitatively similar networks, increasing confidence in model interpretation.

The cpiVAE architecture could be extended towards interesting future directions. One is using the latent space for clustering or subphenotype discovery. If cpiVAE captures biology in its latent variables, those latent variables could be used to cluster individuals or identify functional patterns or associated with disease. This could also reveal subgroups of patients whose proteomic profiles are similar, potentially linking to outcomes or genetics. The cpiVAE latent space itself can even be considered an additional phenotype. Another direction is incorporating uncertainty quantification more formally. The VAE already provides a variance on its outputs through the probabilistic decoder. One could infer confidence intervals for each imputed value and propagate that into downstream analyses, which would be valuable for identifying reliably imputed proteins.

Finally, this study considered cross-imputation between SomaScan and Olink. Given the superior performance of cpiVAE for this applications, a similar architecture could perform well for related applications. Cross-imputation between mass spectrometry and other proteomics methods could be valuable. Mass spectrometry measurements are likely more closely related to absolute concentrations of analytes. Other modalities could also be considered, for example metabolomics data that are often collected for the same cohorts. Imputation within SomaScan or Olink could also be a valuable future application, whether to harmonize different versions of a technology platform or to impute missing data.

In conclusion, cpiVAE advances cross-platform imputation for proteomics, improving the ability to analyze proteomics data within and across cohorts. Performance improves on state-of-the-art methods, whether assessed by technical accuracy, phenotypic or genetic associations, or performance in meta-analysis scenarios. By improving the ability to harmonize data across proteomics cohorts, this method and related deep learning approaches will improved our understanding of the human proteome in health and disease.

## Data and Materials Availability

The cpiVAE source code, trained model weights, preprocessing scripts, and analysis workflows used to generate the results in this manuscript are publicly available under a BSD 2-Clause open-source license at https://github.com/joelbaderlab/cpiVAE_v1 and as an archived release on Zenodo (DOI: 10.5281/zenodo.17245405). These materials include all hyperparameters, model configurations, and evaluation scripts used to reproduce the imputation benchmarks, downstream phenotype association analyses, and interpretability analyses described here. Processed proteomic measurements from the CKB used for model training and internal validation are available from the public FigShare repository https://doi.org/10.6084/m9.figshare.27931350 as described in the original publication. The ARIC proteomics and phenotype data, including SomaScan, Olink, and linked clinical/genetic variables, contain individual-level human participant information and are therefore subject to controlled access. These data can be requested from ARIC Study Coordinating Center under existing data use agreements. Summary-level outputs, including aggregate performance metrics, association statistics, and model evaluation summaries are made available as supplementary files.

## Acknowledgments

This work is supported by National Institute on Aging under award number R01AG085753. MRR was funded in part by NIH/NIDDK grant K01DK141963. The Atherosclerosis Risk in Communities study has been funded in whole or in part with Federal funds from the National Heart, Lung, and Blood Institute, National Institutes of Health, Department of Health, and Human Services, under Contract nos. (75N92022D00001, 75N92022D00002, 75N92022D00003, 75N92022D00004, 75N92022D00005). The authors thank the staff and participants of the ARIC study for their important contributions. SomaLogic Inc. conducted the SomaScan assays in exchange for use of ARIC data. This work was supported in part by NIH/NHLBI grant R01HL134320.

